# *Sleeping Beauty* mutagenesis identifies *BACH2* and other regulators of CD8^+^ T cell exhaustion, persistence *in vivo*, and CAR-T function under tumor-associated chronic antigen stimulation

**DOI:** 10.1101/2025.08.18.670666

**Authors:** Chang-Jung Lee, Alex T. Larsson, Tyler A. Jubenville, Wendy A. Hudson, Carli M. Stewart, Alexander K. Tsai, Adam L. Burrack, Erin E. Nolan, Zach J. Seeman, Yu-Ling Yang, Christopher M. Stehn, Daryl M. Gohl, Margaret Donovan, Nuri A. Temiz, Flavia E. Popescu, Søren Warming, Somasekar Seshagiri, Yun You, Ingunn M. Stromnes, Saad S. Kenderian, David A. Largaespada

**Affiliations:** Department of Genetics, Cell Biology and Development, University of Minnesota, Twin Cities, Minneapolis, Minnesota, USA; Department of Pediatrics, Masonic Cancer Center, University of Minnesota, Minneapolis, Minnesota, USA; T Cell Engineering, Mayo Clinic, Rochester, MN, USA; Mayo Clinic Graduate School of Biomedical Sciences, Mayo Clinic, Rochester, MN, USA; Department of Molecular Pharmacology and Experimental Therapeutics, Mayo Clinic, Rochester, MN, USA; Center for Immunology, University of Minnesota, Minneapolis, MN, USA; Department of Microbiology and Immunology, University of Minnesota, Minneapolis, MN, USA; Department of Medicine, Division of Hematology, Oncology, and Transplantation, University of Minnesota, Minneapolis, MN, USA; University of Minnesota Genomics Center, St. Paul, MN, USA; Institute for Health Informatics, University of Minnesota, Minneapolis, Minnesota, USA; Genentech Inc., San Francisco, California, USA; AntlerA Therapeutics, Toronto, Ontario, Canada; Mouse Genetics Laboratory, University of Minnesota, Minneapolis, Minnesota, USA; Division of Hematology, Department of Medicine, Mayo Clinic, Rochester, MN, USA; Department of Immunology, Mayo Clinic, Rochester, MN, USA

## Abstract

Genes that enhance T cell function represent promising targets for improving engineered T cell therapies for cancer. While extensive CRISPR knockout screens have identified key genes enhancing T cell persistence, employing *Sleeping Beauty* (*SB*) insertional mutagenesis, which induces both gain-(GOF) and loss-of-function (LOF) mutations via the generation of fusion transcripts with endogenous genes, may uncover additional critical factors that previous approaches have overlooked. We developed transgenic mice carrying <u>D</u>oxycycline (Dox)-inducible <u>SB</u> mutag<u>e</u>nesis s<u>y</u>stem (DiSBey) in primary T cells. Using DiSBey, we conducted screens for genetic alterations enhancing T cell persistence under chronic antigen exposure. Specifically, CD8⁺ T cells from Dox-fed DiSBey mice were subjected to repeated anti-CD3 stimulation over 18 days to mimic chronic antigenic stimulation. We then identified *SB* transposon genomic insertion sites and corresponding fusion transcripts from the persistent DiSBey CD8⁺ T cells using enhanced-specificity tagmentation sequencing (esTag-seq) and RNA-seq, respectively. Under chronic stimulation, *SB*-mutagenized CD8⁺ T cells exhibited improved persistence and reduced terminal exhaustion phenotype. Across six independent screens, we identified 38 genes that were recurrently targeted by the *SB* transposon T2/Onc2 and differentially expressed under chronic anti-CD3 stimulation stress. Among these, T2/Onc2 insertions into *Bach2* and *Elmo1* were repeatedly found at the genomic level and were associated with altered nascent transcript expression. *Bach2*, known as a key regulator of T cell memory formation and resistance to chronic viral infection but less characterized in engineered T cells for cancer therapy, was found to enhance *in vivo* tumor persistence in the B16-Ova tumor model. We showed that ectopic *Bach2* expression levels influence engineered T cell differentiation lineage. A Bach2^low^ signature allowed differentiation into both KLRG1⁺ and CD62L⁺ phenotypes, whereas Bach2^high^ restricted differentiation predominantly to the CD62L⁺ subset. Finally, in human CART19-28ζ cells, *BACH2* overexpression enhanced cytotoxicity and improved tumor control following chronic cancer stimulation. Controllable *SB* mutagenesis using DiSBey mice provides a novel platform for functional screening of genes that improve T cell therapeutic phenotypes. Our findings highlight a dose-dependent role of BACH2 in enhancing the function of engineered T cells under conditions of chronic antigenic stimulation.

## Background

Immunotherapies based on cytotoxic T cell activity, such as chimeric antigen receptor (CAR)-T cell therapy, have been successful for certain hematological and solid tumors, but the initial responsiveness and the durability of responses have been major problems^1,2^. These problems are, in part, associated with T cell exhaustion^3^, characterized by a progressive hypofunctional state of cancer-reactive T cells experiencing chronic stimulation on T cell receptor (TCR) signaling in the tumor microenvironment (TME)^3^. The exhaustion phenotype is characterized by the increasing expression of terminally exhausted immunoinhibitory receptors, reduced *in vivo* expansion, reduced effector function, and increased apoptosis^4–9^. T cell exhaustion has been directly associated with the failure of checkpoint blockade, CAR-T cell, and transgenic TCR-T cell therapies^3,7,9^.

*Sleeping Beauty* (*SB*) mutagenesis, by combining the activation of *SB* transposase to mobilize a mutagenic *SB* transposon T2/Onc2 into the genome, is a powerful forward genetic screening tool to identify oncogenic drivers^15–17^. While *SB* was used to identify oncogenic drivers for T cell acute lymphoblastic leukemia^18^ and endogenous T cell trafficking mechanisms to tumors^19^, it has not been utilized for identifying genes to enhance engineered T cells due to its pre-leukemic risk when *SB* mutagenesis is active continuously over many weeks. The T2/Onc2 construct includes an MSCV promoter, a splice donor, and a bidirectional polyadenylation signal, all flanked by splice acceptors (Figure 1B)^16,21^. Depending on the integration site, T2/Onc2 insertion can result in both gain-of-function (GOF) and loss-of-function (LOF) mutations. These structural elements facilitate the generation of truncated structural variants, which further increase the diversity of the resulting mutant pool. In addition, not only can the DNA insertion sites of T2/Onc2 be mapped at the genomic level, the endogenous gene transcripts activated or truncated by the insertion of T2/Onc2 can also be detected using RNA sequencing (RNA-seq)^22^. These features allow *SB* mutagenesis to uncover critical genetic mechanisms that may be overlooked by CRISPR-based approaches.

**Figure 1.**
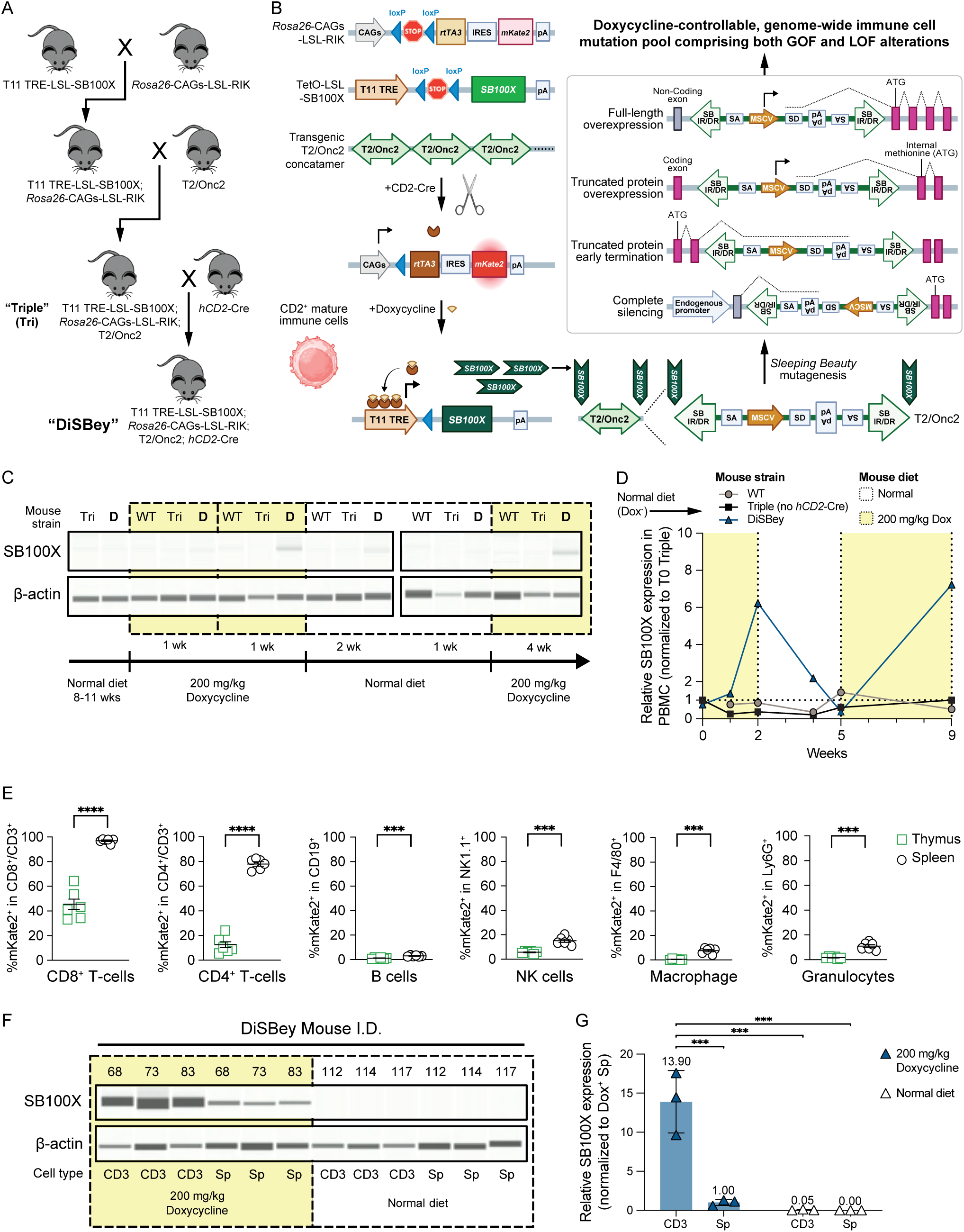
DiSBey mouse model allowed doxycycline-controllable *Sleeping Beauty* (*SB*) mutagenesis in T cells. A) Breeding scheme to generate DiSBey mice. B) Schematic of *SB* mutagenesis for DiSBey mice. C) SB100X and β-actin expression in the PBMCs collected from WT, triple transgenic (lack *hCD2*-Cre, “Tri”), or DiSBey (“D”) mice with intermittent Dox treatments. D) Relative SB100X expression in (C). SB100X expression was normalized by β-actin expression and time 0 (T0) triple mouse. E) Cre-recombination efficiency in immune cell compartments in the thymus (green) and the spleen (blue). Cre-activated fluorescent mKate2 was used as a reporter. n= 6 mice. F) SB100X expression in T cells (CD3) or splenocytes (Sp) from DiSBey mice treated with or without Dox-containing diet. G) Relative SB100X expression in (F). SB100X expression was normalized by β-actin expression and DiSBey splenocytes. (Two-way ANOVA with Tukey‘s test, n=3 mice per group)

Here, we created a novel transgenic mouse model, DiSBey, allowing controllable *SB* mutagenesis in primary T cells. CD8^+^ T cells from the DiSBey mice were selected by anti-CD3 chronic stimulation, and the mutagenic *SB* transposon T2/Onc2 DNA insertion sites and nascent mRNA fusion transcripts were identified by next-generation sequencing. From six independent DiSBey screens, 38 genes were repeatedly found to be mutagenized by *SB* under chronic antigenic stimulation, with *Bach2* and *Elmo1* both detected at the genomic and transcript levels. *Bach2*, a master regulator for T cell stemness and memory phenotype^23–26^, was found to be strongly enriched in the chronic-stimulation-resistant DiSBey CD8^+^ T cells. Ectopic Bach2 OE improved the persistence of CD8^+^ T cells in both *in vitro* and *in vivo* in B16-Ova models. OE of BACH2 in human CART19-28ζ also improved targeted cytotoxicity and therapeutic outcome after chronic stimulation. Further, higher BACH2 expression correlated with the responsiveness of pembrolizumab and ipilimumab in melanoma patients. Taken together, pioneering the use of *SB* mutagenesis to improve engineered primary T cells, we identified the crucial role of BACH2 in sustaining CD8^+^ T cell functionalities for cancer immunotherapies.

## Methods

### DiSBey mouse models generation

C2 mouse embryonic stem cells (C57BL/6N) were electroporated with linearized T11-LSL-SB100X plasmid (**Supplementary Figure 1A**, MGI allele 8214726), and clones were selected using established methods. Resulting clones were screened by long-range PCR followed by sequence confirmation and correctly targeted T11-LSL-SB100X clones were injected into albino blastocysts followed by implantation into pseudopregnant females. High percentage chimeric male offspring were crossed to albino C57BL/6N females to generate N1 T11-LSL-SB100X heterozygous mice. Subsequent N1 mice were backcrossed to C57BL/6J mice for at least four generations to create congenic mice B6J.B6N-Col1a1^tm1(tetO-sb100x)Dla^ (MGI strain 8214988). SB100X mice were then crossed to mice carrying the Rosa-CAGS-LSL-rtTA3-IRES-mKate2 allele^28^ [Gt(ROSA)26Sor^tm2(CAG-rtTA3,-mKate2)Slowe^], hCD2-Cre^29^ (C57BL/6-Tg(CD2-cre)1Lov/J), and T2/Onc2^17^ [TgTn(sb-T2/Onc2)6113Njen for DiSBey^Chr1^ or TgTn(sb-T2/Onc2)6070Njen for DiSBey^Chr4^] to create DiSBey mice. Genotyping PCR primer sequences are listed in **Supplementary Table 1**.

### Flow cytometry

CytoFLEX S or BD FACSAria II was used for flow cytometry analysis or sorting. Flow cytometry data were analyzed by FlowJo™ v10 or Kaluza 2.3. All antibodies used in this study are listed in **Supplementary Table 2**.

For intracellular cytokine staining of adoptively transferred mouse T cells, single-cell suspensions isolated from different mouse organs were treated with Brefeldin A (BD 555029) and PMA/Ionomycin (BioLegend 423301) or 1 uM SIINFEKL peptide (Anaspec AS-60193-1) for 4 hours. Cells were washed, stained with fixable viability dyes, and antibodies against surface markers indicated above. After surface staining, cells were fixed and permeabilized before staining with antibodies against intracellular cytokines indicated above.

For intracellular cytokine staining of human CAR-T cells, CAR-T cells were co-cultured with JeKo-1 cells at a 1:5 ratio for 4 hours after adding 2 µL CD28 (BD 348040), and 1 µL monesin (BioLegend 420701) per 200µL. Cells were then stained with viability dye, fixed, and permeabilized before staining.

### In vitro chronic TCR stimulation

Adapted from Belk et al.^13^. After anti-CD3/anti-CD28 activation, murine CD8^+^ T cells were seeded in cultured plates coated with (for chronic stimulation) or without (for acute stimulation) 5 µg/ml anti-CD3ε (BioLegend 100359). Afterwards, 75% of TCM in each well was changed every odd day, and T cells were replated at 10^6^ cells per ml every even day until day 18.

### In vitro CAR-T chronic exhaustion assay

To evaluate the effect of chronic stimulation on BACH2 OE CAR-T cells, we used an *in vitro* exhaustion model as previously described^33^. On day 8 of CAR-T production, cells were co-cultured with JeKo-1 tumor cells at a 1:1 ratio. Restimulation with the same number of JeKo-1 cells was performed on days 10, 12, and 14. On day 15, CAR-T cells were isolated using CD4 and CD8 positive selection beads (Miltenyi Biotec 130-045-101 and 130-045-201), and a subset was subjected to a second week of chronic stimulation. These cells were co-cultured with JeKo-1 cells at a 1:1 ratio and restimulated on days 17, 19, and 21. Final isolation was performed on day 22. CAR-T cells from days 15 and 22 were then used in functional assays to assess changes in cell activity.

### Statistical analysis

Statistical methods and the number of samples tested (n) for each experiment can be found in the figure legends. Unless otherwise specified, statistical significance was determined as follows: ns, p > 0.05; *, p < 0.05; **, p < 0.01; ***, p < 0.001; ****, p < 0.0001.

Detailed information on cell lines, mouse strains, tissue processing, T cell culturing and assays, and sequencing data processing and analysis can be found in **Supplementary Methods**.

## Results

### Generation of transgenic mouse models allowing controllable *SB* mutagenesis in T cells

To generate T cell mutant pools for phenotype screening beneficial to ACT while minimizing leukemogenic risk, we developed a temporally controllable SB100X transposase transgene. Specifically, we created the congenic mouse model B6J.B6N-Col1a1^tm1(tetO-sb100x)Dla^, carrying a loxP-stop-loxP–flanked SB100X transgene driven by the tetracycline-responsive element (TRE) knocked into the *Col1a1* locus (Supplementary Figure 1A). This line was then bred with Rosa26-LSL-rtTA3-mKate2^28^, hCD2-Cre^29^, and T2/Onc2^17^ mice to create a line with <u>D</u>ox-inducible *<u>SB</u>* mutag<u>e</u>nesis s<u>y</u>stem (DiSBey) in hCD2-active cells (Figure 1A, 1B). To expand mutagenesis pools and control for local hopping, we used two T2/Onc2 strains with concatemer insertion sites on chromosomes 1 and 4 (TgTn(sb-T2/Onc2)6113Njen and TgTn(sb-T2/Onc2)6070Njen^17^), resulting in DiSBey^Chr1^ and DiSBey^Chr4^ lines.

### *In vivo* SB100X activity is regulated in T cells from DiSBey mice

SB100X expression was inducible only in DiSBey mice via Dox-containing diets and absent in C57BL/6J wild-type (WT) or triple transgenic mice (DiSBey littermates lacking hCD2-Cre transgene) (Figure 1C, 1D). Cre-mediated recombination was efficient in CD8^+^ and CD4^+^ T cells, low in NK cells and macrophages, and minimal in B cells and granulocytes in DiSBey splenocytes. In the thymus, only CD8^+^, CD4^+^ T cells, and NK cells showed more than 5% of recombination (Figure 1E). *SB* mutagenesis in T cells was also dox-dependent both *in vitro* and *in vivo* (Figure 1F, 1G, Supplementary Figure 1B, 1C).

### *SB* mutagenesis enhances CD8^+^ T persistence under chronic TCR stimulation

To identify *SB*-mutagenized CD8⁺ T cell clones with enhanced persistence under chronic TCR stimulation, we employed an *in vitro* chronic exhaustion assay^13^. In this model, polyclonal murine CD8⁺ T cells were subjected to repeated stimulation with anti-CD3 antibodies to simulate prolonged TCR signaling and induce key features of T cell exhaustion induced *in vivo*^13^. CD8⁺ T cells from DiSBey and WT mice (on Dox diet for six weeks) were activated and cultured under either chronic (anti-CD3-coated plates + IL-2) or acute (uncoated plates + IL-2) stimulation. Cells were replated every two days and harvested for analysis until day 18 (Figure 2A). Chronic stimulation reduced viability and expansion, but DiSBey T cells showed a significant viability advantage over WT from day 14 onward (Figure 2B). No expansion difference was observed (Supplementary Figure 1D). By day 18, DiSBey^Chr4^ T cells showed a shift from PD-1⁺/TIM-3⁺ to PD-1⁺/TIM-3⁻ phenotype (Figure 2C, 2D), not seen in DiSBey^Chr1^. This suggests SB mutagenesis enriches for CD8⁺ T cells with enhanced persistence and reduced exhaustion.

**Figure 2.**
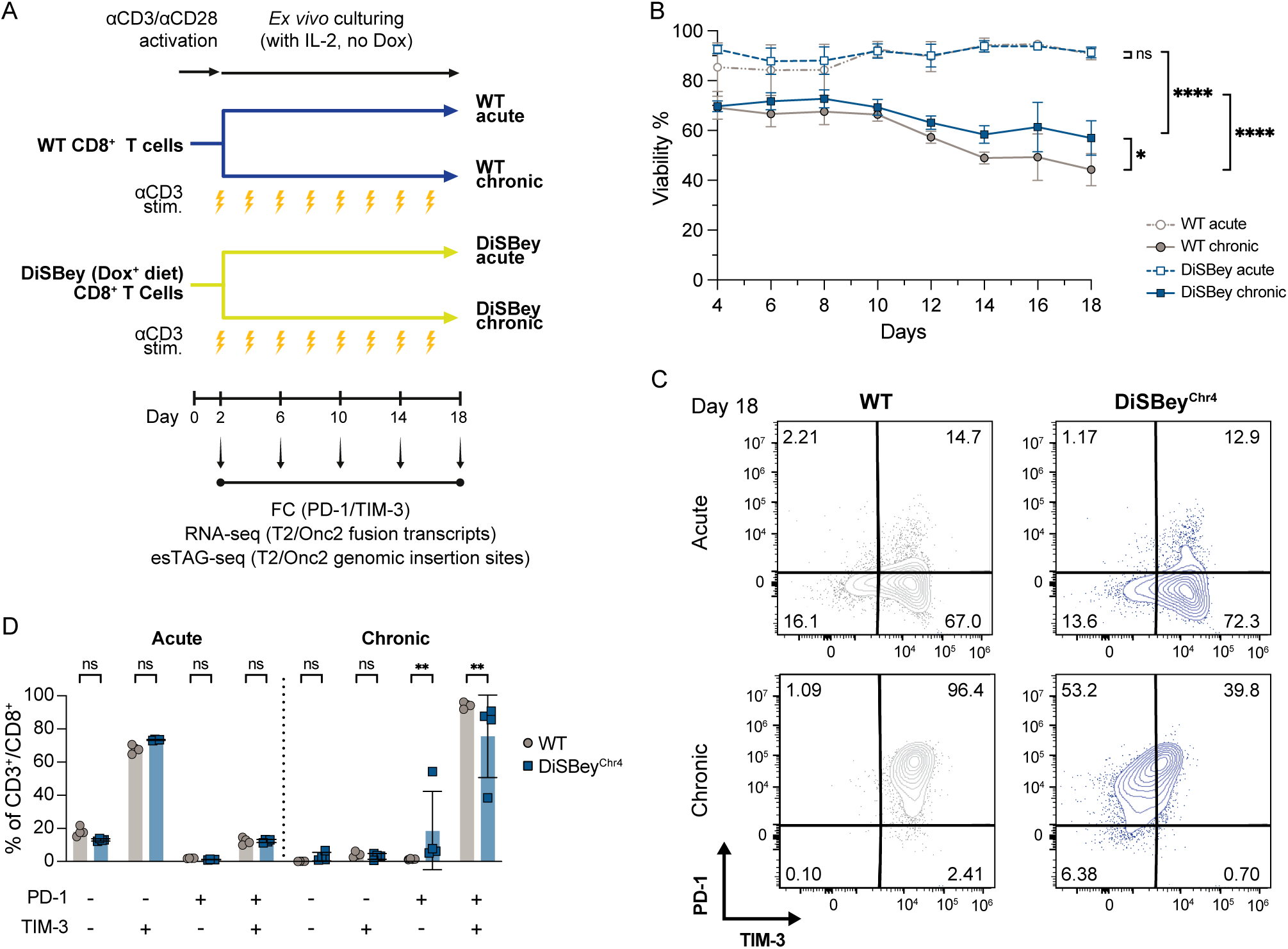
*SB-mutagenized* CD8^+^ T cells from DiSBey mice exhibited enhanced persistence under chronic TCR stimulation. A) Schematic of chronic TCR stimulation screen using DiSBey. B) Viability of cultured cells undergoing chronic or acute stimulation *in vitro.* (Two-way ANOVA with Tukey’s test, n=7 for WT and 6 for DiSBey mice from 2 independent experiments) C) Representative figures for exhaustion markers PD-1 and TIM-3 from WT and DiSBey^Chr4^ at day 18 of chronic TCR stimulation. Results were gated at the CD3^+^/CD8^+^ population. D) Quantification of PD-1 and TIM-3 population from WT and DiSBey^Chr4^ at day 18 of chronic TCR stimulation. (Two-way ANOVA with Tukey‘s test, n=4 per group)

### *SB* genomic insertion and fusion mRNA transcript profiles identify unique and recurrent insertion mutations in the persisting DiSBey CD8^+^ T cells

To identify the genes driving persistence under chronic TCR stimulation, we performed enhanced-specificity tagmentation-assisted sequencing (esTag-seq) to map genomic T2/Onc2 insertions and RNA-seq to detect resulting fusion transcripts. Thousands of insertion events were identified and were prioritized if (1) differentially present under chronic vs. acute conditions, or (2) uniquely present in DiSBey samples (Figure 3A, 3B). Insertions to *Bach2*, a known transcription repressor associated with T cell stemness, memory formation, and resistance to chronic viral infection^23–26^ but less understood in engineered T cells against tumors, emerged as a top hit with both enriched genomic insertions and fusion transcripts (Figure 3B, 3C). Additional genes, such as *Bcl2*, *Akt2*, *Nabp1*, and *Myh9*, showed evidence of transcriptional perturbation or genomic insertion, indicating other candidates that potentially drive T cell persistence. Sense T2/Onc2 Insertions upstream of *Bach2* coding sequences indicated full-length transcriptional activation (Figure 3D). Bach2 levels were restored in chronically stimulated DiSBey T cells but remained low in WT (Figure 3E). In contrast, *Elmo1* insertions were antisense, likely leading to repression (Supplementary 2A, 2B).

**Figure 3.**
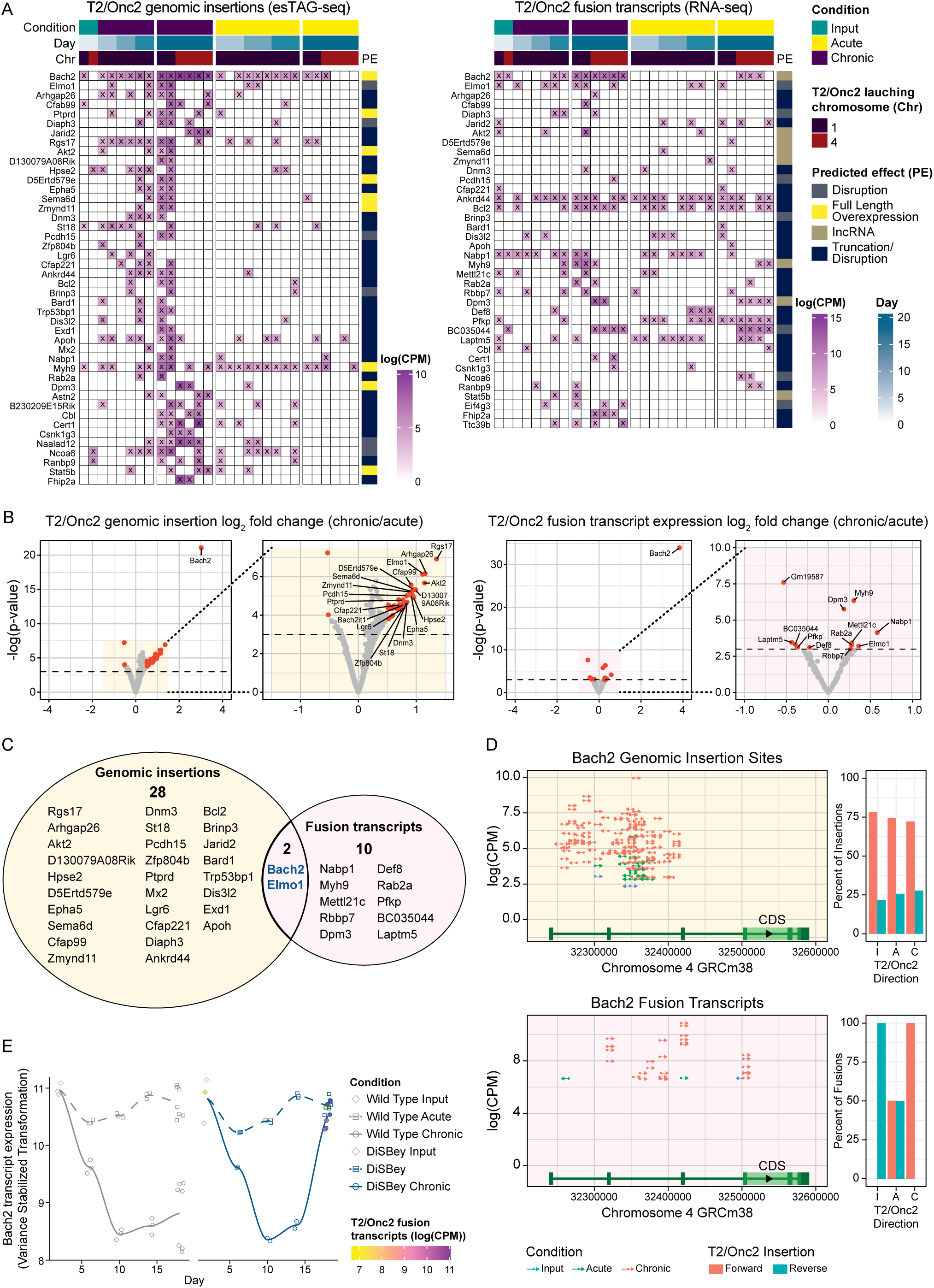
Interrogation of genomic insertion and nascent fusion transcripts of T2/Onc2 in DiSBey CD8^+^ T cells underwent chronic TCR stimulation. A) Genes identified with differentially expressed or exclusively detected T2/Onc2 insertions. B) Volcano plots showing differential T2/Onc2 genomic insertions (left) and fusion transcripts (right) between day 18 acute and chronic stimulated DiSBey CD8^+^ T cells. C) Venn diagram of differential T2/Onc2 genomic insertions and fusion transcripts. D) T2/Onc2 genomic (upper) and transcriptomic (lower) insertion diversity in *Bach2*. E) Normalized *Bach2* expression from DESeq2 and compared against the counts per million T2/Onc2 fusion *Bach2* transcripts over *in vitro* chronic stimulation assay.

### Ectopic Bach2 OE maintains CD8^+^ T cell persistence against chronic stimulation in a dose-dependent manner

We next examined how Bach2 supports CD8⁺ T cell persistence under chronic stimulation and whether its overexpression affects anti-tumor cytotoxicity. OT-I CD8⁺ T cells were transduced with Bach2-IRES-GFP (Bach2 OE) or control vector (GFP) (Supplementary 3A) and then subjected to chronic anti-CD3 stimulation (Figure 4A). Bach2 OE caused enhanced viability, reduced apoptosis, and similar cytotoxicity to that of control cells after 18 days of chronic stimulation (Figures 4B-4F). Interestingly, the enhanced persistence due to Bach2 OE appeared to be dose-dependent, as low Bach2 OE (Bach2^low^) had preferential enrichment during chronic TCR stimulation compared with high Bach2 OE (Bach2^high^) (Figure 4G, 4H, Supplementary Figure 3D). Meanwhile, proliferation was associated with the level of ectopic Bach2 expression, as Bach2^high^ populations showed the poorest proliferation, and Bach2^low^ populations exhibited intermediate proliferation compared to their control counterparts (Figure 4G, 4I). This indicates that ectopic Bach2 expression level tunes T cell behavior during chronic antigen exposure.

**Figure 4.**
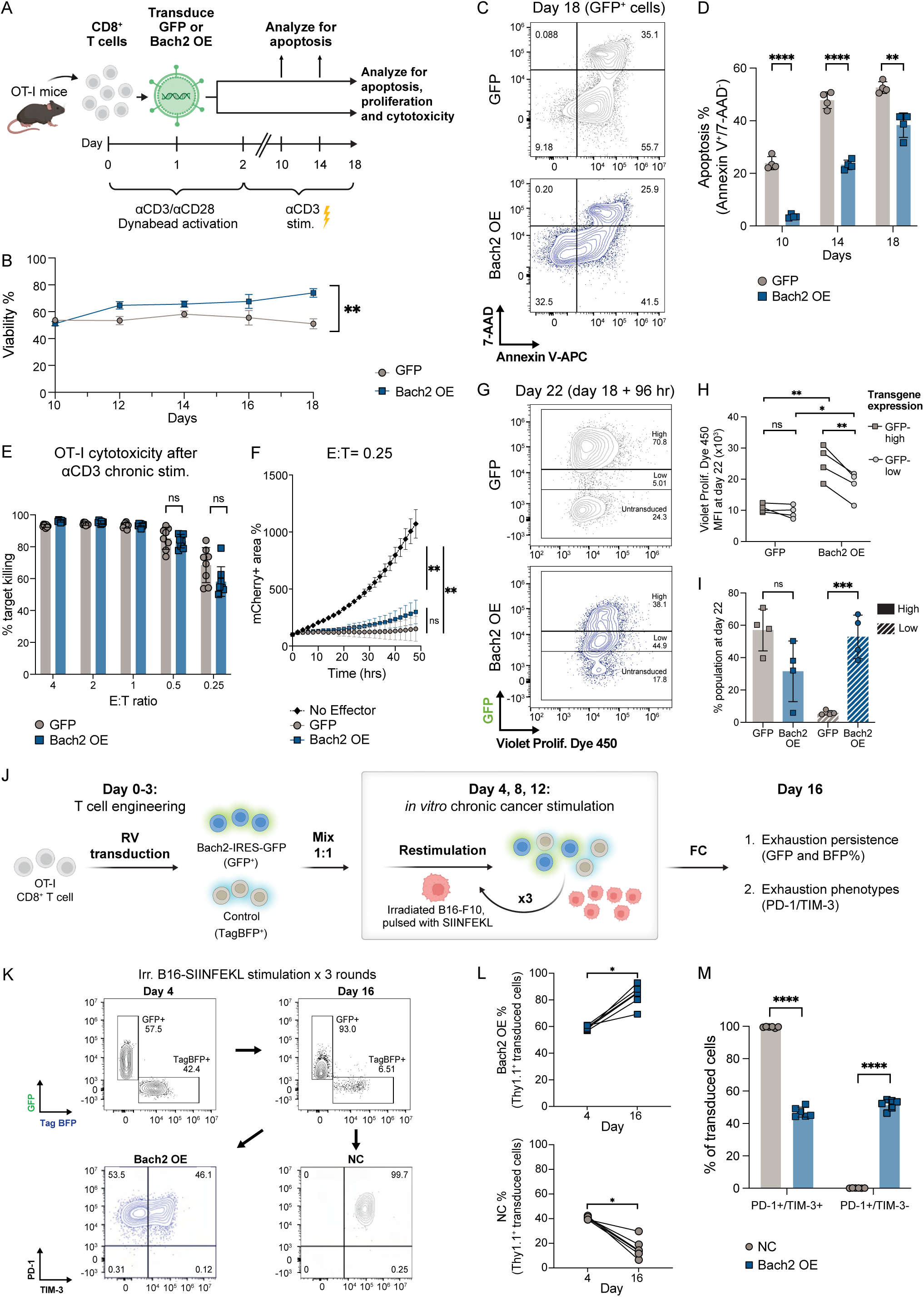
Bach2 OE promoted enhanced persistence of CD8^+^ T cells under both chronic TCR and chronic cancer stimulations. A) Schematic for functional validation of Bach2 OE CD8^+^ T cell after chronic anti-CD3 stimulation for panels B to I. n=4 mice. B) Viability of OT-I T cells transduced with GFP or Bach2-GFP under chronic TCR stimulation with anti-CD3-coated plates (Two-way ANOVA with Tukey‘s test). C) Representative figures for 7-AAD/Annexin V apoptosis analysis of day 18 chronically stimulated OT-I cells. Results were gated at the transduced (GFP^+^) population. D) Quantification of apoptotic OT-I T cells at each indicated day of chronic TCR stimulation. (unpaired t-test). E and F) Cytotoxicity assessment of OT-I co-cultured with JW18-SIINFEKL for 48 hours. E, day 18 transduced OT-I co-cultured at various E:T ratios. F, Time-lapse of mCherry^+^ areas for E:T ratio=0.25. (unpaired t-test, n=4 mice× 2 technical repeats per group). G) Representative figures for proliferation analysis of day 18 chronically stimulated OT-I. Results were gated on the live cells. H) MFI of Violet Proliferation Dye 450 in (F). (Paired t-test for high/low comparison, unpaired t-test for GFP/Bach2 OE comparison) I) Population frequency based on GFP intensity in (F). (Paired t-test) J) Schematic for *in vitro* competition assay for persistence under chronic cancer cell stimulation. K) Representative figures depicting GFP^+^ Bach2-OE and TagBFP^+^ NC population rate, and PD-1 and TIM-3 expression for the respective populations for day 4 and 16. Irr.B16-SIINFEKL: irradiated-B16-F10 pulsed with SIINFEKL. Results were gated at Thy1.1^+^ transduced cells. L and M) Quantification of replicates from day 16 samples in (K). L, Fluorescent population frequency. M, PD-1 and TIM-3 expression for the respective populations (Paired t-test for fluorescent population comparison, unpaired t-test for exhaustion phenotype comparison, n=3 biological repeats × 2 technical repeats per group)

Next, we performed an *in vitro* competition assay to assess Bach2 OE under chronic antigenic stimulation from cancer cells. OT-I-PL (Thy1.1⁺) CD8⁺ T cells were transduced with either Bach2 OE or a negative control vector expressing TagBFP, and mixed at a 1:1 ratio before repeatedly stimulated with irradiated B16-F10 melanoma cells pulsed with SIINFEKL peptide for three rounds (Figure 4J). After three rounds of stimulation, the Bach2 OE population significantly outcompeted the TagBFP population, indicating enhanced persistence (Figure 4K, 4L). Further, these Bach2 OE cells were enriched for a PD-1⁺/TIM-3⁻ population, while most NC cells displayed a PD-1⁺/TIM-3⁺ terminally exhausted phenotype (Figure 4K, 4M). These results show Bach2 OE promotes CD8⁺ T cell persistence and reduces exhaustion under chronic tumor engagement.

### Bach2 OE increases *in vivo* T cell persistence without improving tumor control

We next examined whether Bach2 OE could enhance persistence *in vivo* within an immune-competent background. OT-I-PL CD8^+^ T cells transduced with either Bach2 OE or TagBFP control were mixed and transferred into B16-Ova-bearing C57BL/6J mice (Figure 5A). Bach2 OE OT-I cells in tumors *in vivo* also showed a selective advantage at day 14, but not at earlier time points (Figure 5B). Similarly, we observed that Bach2 OE OT-I cells demonstrated a remarkably enhanced ability to persist in the draining lymph nodes, spleens, and circulating blood (Figure 5B). However, despite improved persistence, no survival benefit was observed (Supplementary Figure 4), suggesting the improved persistence by Bach2 OE is insufficient for tumor control.

**Figure 5.**
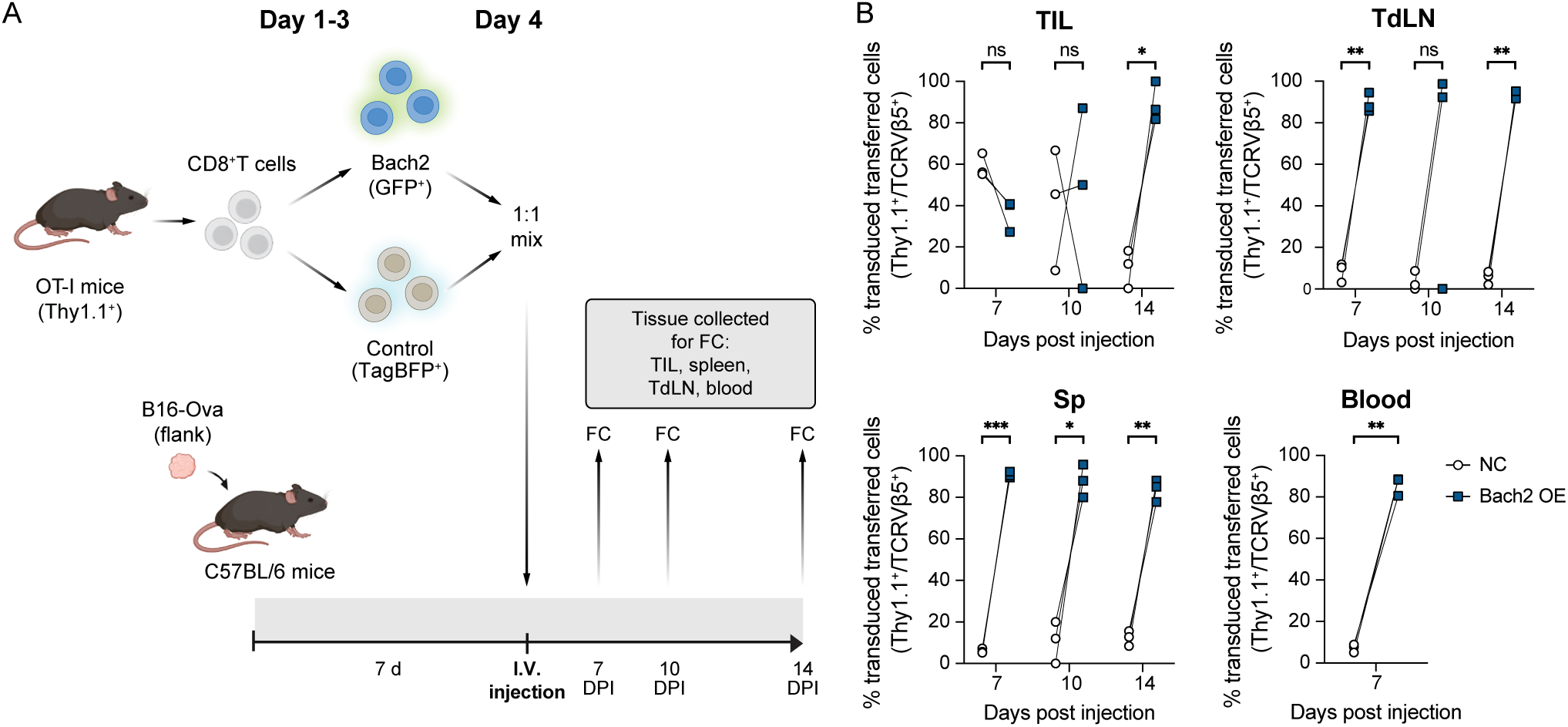
Bach2 OE improved transferred T cell persistence against B16-Ova *in vivo*. A) Schematic for *in vivo* competition assay for transferred T cell persistence in C57BL/6J B16-Ova model. B) Population rate of Bach2 (GFP^+^) and control (TagBFP^+^) in transferred OT-I T cells isolated from each indicated time point and organs. TIL: tumor-infiltrating lymphocytes. TdLN: tumor draining lymph nodes. Sp: spleens. Results were gated at live Thy1.1^+^/TCRVβ5^+^ fluorescent cells. (unpaired t-test, n=3 mice per group)

### Low-level ectopic Bach2 OE enables both effector and memory phenotypes

To investigate the function of T cells ectopically Bach2 OE in a second well-characterized tumor, we employed an orthotopic pancreatic ductal adenocarcinoma (PDA) model^37,38^. An Ova^+^ PDA cell line^31^ was surgically re-implanted into the pancreas of C57BL/6J mice, followed by transferring Thy1.1^+^ OT-I T cells transduced with Bach2 OE or GFP control vectors (Figure 6A). Cognate SIINFEKL peptide restimulation of transferred OT-I T cells demonstrated that Bach2 OE OT-I cells exhibited reduced production of IFN-γ and TNF-α compared to controls in all organs collected. However, upon PMA and ionomycin restimulation, Bach2 OE T cells displayed cytokine production capacities for IFN-γ and TNF-α comparable to controls, while IL-2 production was notably elevated in cells isolated from both tumors and draining lymph nodes relative to the control counterparts (Figure 6B). These results indicate that Bach2 OE allows CD8^+^ T cells to maintain high secretion capacity of IFN-γ and TNF-α and even enhances IL-2 secretion, but it also reduces the sensitivity of these cells to TCR signaling, which in turn impairs their overall effector function. The Bach2 OE group also showed a reduced proportion of KLRG1^+^/CD62L^-^ cells (short-lived effector T cells with higher cytotoxic potential)^23,39^ and increased proportion of KLRG1^-^/CD62L^+^ cells (long-lived central memory T cells with reduced cytotoxicity) ^23,39^ in the spleen, compared to control T cells (Figure 6C, 6D). Interestingly, we noted an enrichment of Bach2^low^ T cells as indicated by lower GFP levels compared to input (Figure 6E). These Bach2^low^ cells consisted of both KLRG1^+^/CD62L^-^ or KLRG1^-^/CD62L^+^ subsets. In contrast, Bach2^high^ cells predominantly assumed a KLRG1^-^/CD62L^+^ phenotype, whereas control cells tended to adopt a KLRG1^+^/CD62L^-^ phenotype (Figure 6E, 6F). Together, these findings indicate that while Bach2 OE promotes memory-like phenotypes in T cells, it also attenuates TCR signaling and limits effector functions. Notably, this effect depends on the ectopic Bach2 expression level, as lower levels of ectopic Bach2 OE permit phenotypic flexibility between effector and memory states, whereas high levels enforce a fixed memory-like state.

**Figure 6.**
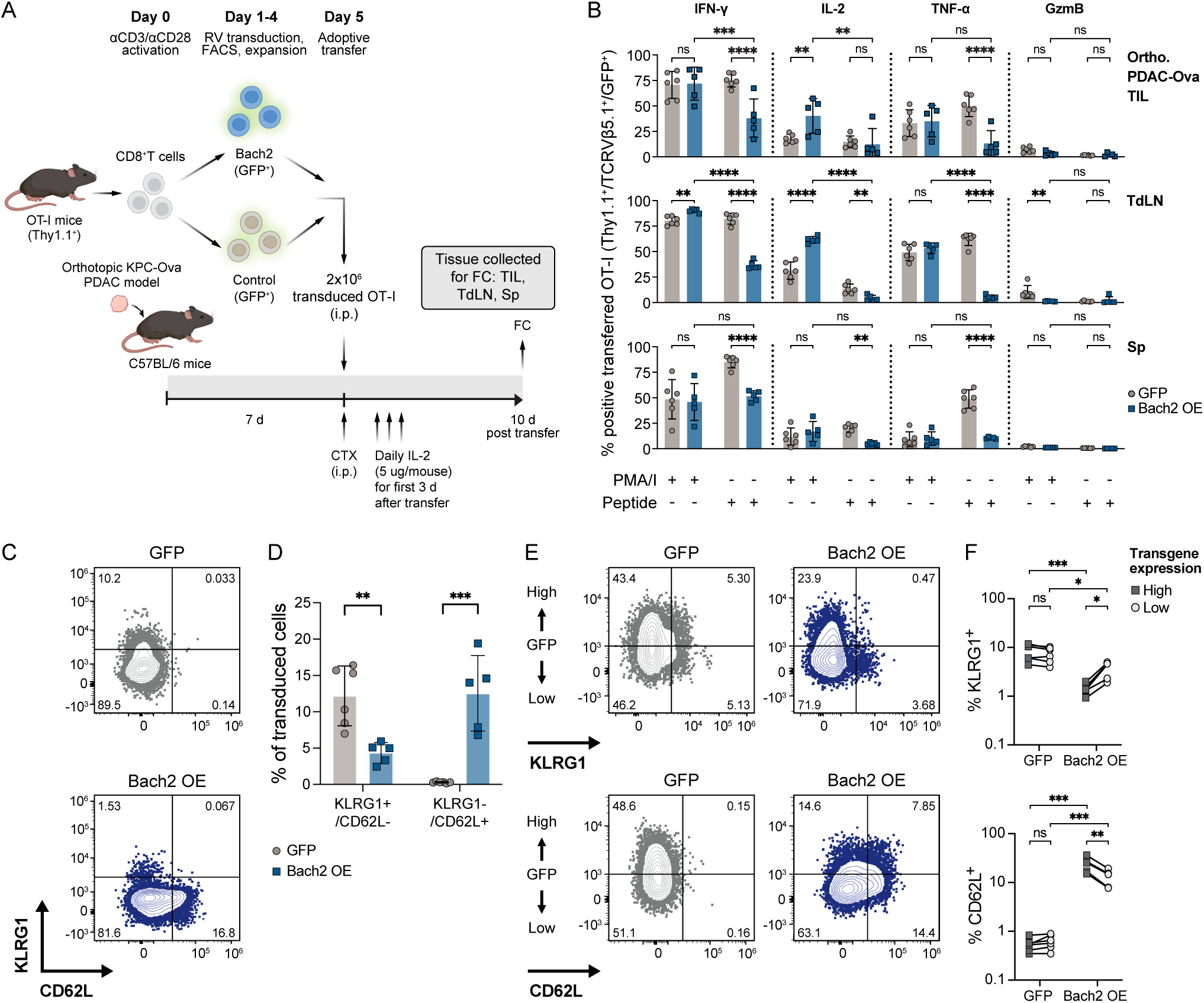
Bach2 OE decreased TCR signaling sensitivity and retained lineage capacity to both KLRG1^+^ and CD62L^+^ lineages. A) Schematic of assessing transferred T cell phenotypes in orthotopic PDAC-ova model. n=6 mice injected with OT-I GFP, and 5 mice injected with OT-I Bach2 OE for panels B to F. B) Cytokine secretion following PMA/I or SIINFEKL peptide stimulation for transferred cells isolated from the indicated organ. TIL: tumor-infiltrating lymphocytes. TdLN: tumor draining (inguinal) lymph nodes. Sp: spleens. Results were gated at live Thy1.1^+^/TCRVβ5^+^/GFP^+^ cells. (Two-way ANOVA with Tukey’s test) C and D) Representative figures (C) and frequency (D) of KLRG1^+^/CD62L^-^ or KLRG1^-^/CD62L^+^ in transferred OT-I isolated from spleens of mice carrying orthotopic PDAC-Ova tumors (unpaired t-test). E) Representative figures for transgene expression levels (GFP expression as a proxy) with KLRG1 (upper) and CD62L (lower) from (D). F) Quantification of population frequency of KLRG1^+^ (upper) or CD62L^+^ (lower) rate from GFP-high or -low population from (E). (Unpaired t-test for comparisons between GFP or Bach2 OE group, paired t-test for comparisons within GFP or Bach2 OE group).

### BACH2 OE in human CART cells contributed to improved survival

We next investigated whether OE of human BACH2 enhances the therapeutic performance of CAR-T cells under conditions of chronic stimulation. To this end, human CART19-28ζ cells were transduced with BACH2 OE or GFP-only control vector (Supplementary Figure 5A, 5B). These engineered CAR-T cells were then subjected to an *in vitro* chronic stimulation protocol, wherein CD19^+^ Jeko-1 target cells were replenished every other day for 14 days post-CAR-T manufacturing^33^ (Figure 7A). Surprisingly, BACH2 OE CART19-28ζ cells showed a trend towards an improved persistence for the first 7 days of chronic stimulation (p=0.1004), but not at the end of the assay (Supplementary Figure 5D). Checkpoint and cytokine showed interindividual variation between donors (Supplementary Figure 5E, 5F). However, following chronic stimulation with Jeko-1 cells, BACH2 OE CART19-28ζ cells exhibited enhanced cytotoxicity *in vitro* and improved tumor control *in vivo* (Figure 7B-7E). Next, we analyzed the association between BACH2 expression and treatment responses in patients undergoing CAR-T therapy. BACH2 expression was elevated in CD8^+^ T cells of CAR-T responders in the TanCAR7 T trial^40^ (Figure 7F), with a similar trend in CD4^+^ CAR-T (p=0.06) and naïve subsets (p=0.09) and in the ZUMA-1 trial^41^ (p=0.15) (Supplementary Figure 6A).

**Figure 7.**
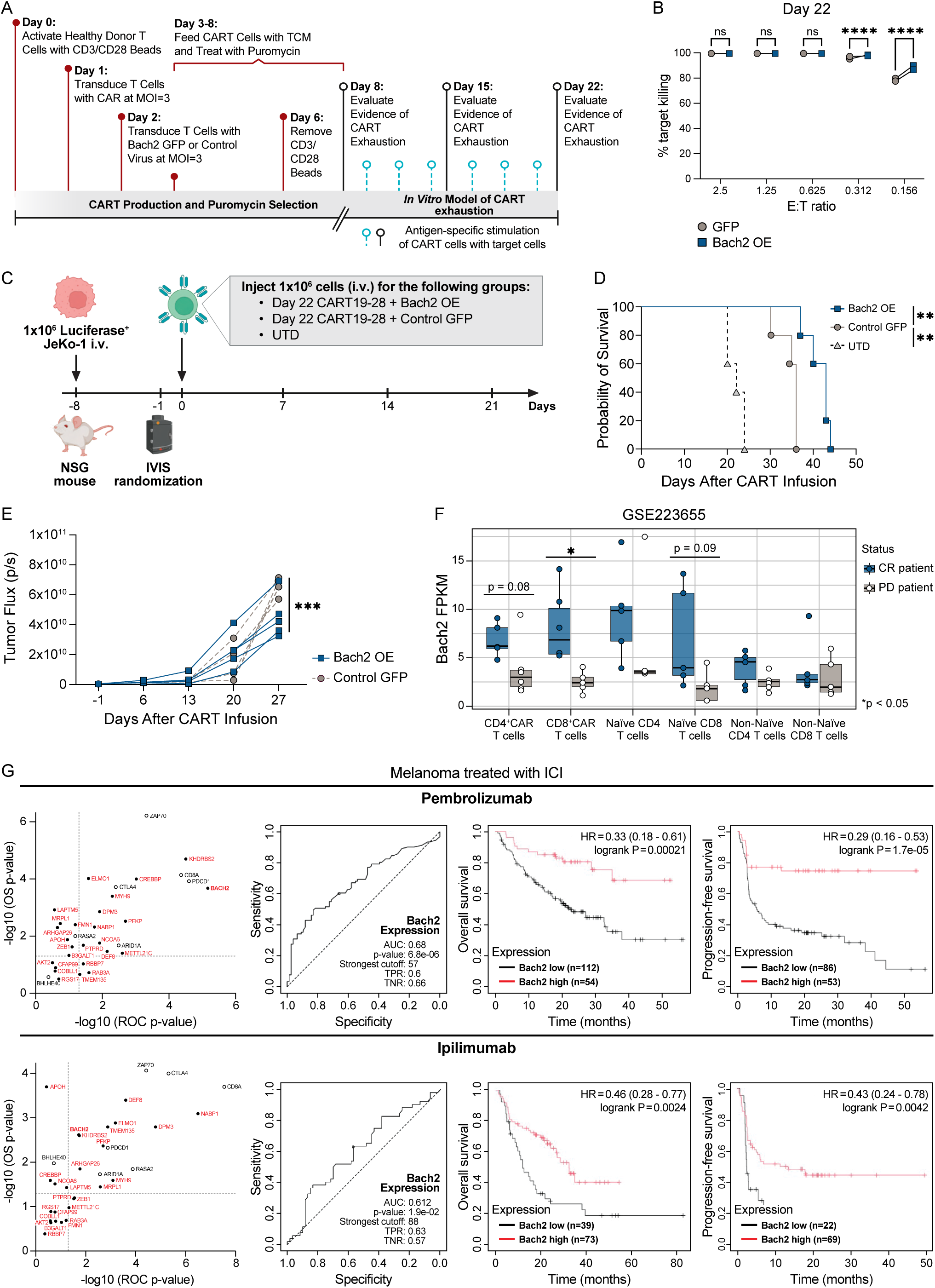
BACH2 could represent a novel target for T cell therapies. A) Schematic of evaluating BACH2 OE human CAR19-28ζ with an *in vitro* chronic exhaustion assay. B) Cytotoxicity assessment of BACH2 OE human CAR19-28ζ after chronic exhaustion assay. Chronically exhausted CAR-T cells were co-cultured with Jeko-1 (Luciferase^+^) at various E:T ratios for 48 hours before measuring luciferase activity. (Paired t-test, n=3 donors per group) C) Schematic of evaluating the tumor control of chronically exhausted BACH2 OE human CAR19-28ζ. n=5 mice per group from panel C-E. D) Overall survival curve based on JeKo-1 xenograft mouse model comparing treatment with Day 22 control GFP or BACH2 OE CART19-28ζ cells. (Log-rank (Mantel–Cox) test) E) Bioluminescence of the tumor growth in the JeKo-1 xenograft mouse model before the first death at day 27. F) Comparison of BACH2 expression level with treatment responsiveness of CAR-T products in the TanCAR7 T trial. CR: complete response; PD, progressive disease. (Unpaired t-test, n=6 patients per group) G) Correlation analysis of genes identified from the DiSBey chronic TCR stimulation screens (Figure 3B) expression level with melanoma patients treated with ICI^42^. For the left panels, genes identified from this study were labeled in red, while other previously identified genes were labeled in black.

### Exhaustion resistance genes identified by SB mutagenesis may influence response to immune checkpoint inhibition

We assessed the relationship between the expression of candidate genes from our exhaustion screen and clinical outcomes in melanoma patients treated with Pembrolizumab or Ipilimumab using a curated clinical database of human immune checkpoint inhibitor (ICI) trials^42^. Among candidates identified from the DiSBey screen, *BACH2, ELMO1, NABP1, PFKP, MYH9, DEF8,* and *DPM3* expression correlated with responsiveness and survival for both ICI therapies (Figure 7G, Supplementary Figure 6B). However, these gene expressions measured from bulk RNA may not directly reflect their expression in T cells *in situ*.

## Discussion

The lack of *in vivo* persistence, especially under chronic antigenic stimulation, is widely considered an obstacle to successful T cell-based immunotherapies against cancers^1,2^. We present DiSBey mice as a novel screening model with Dox-regulated *SB* mutagenesis to identify genes that improve T cell persistence under repeated TCR stimulation. Here, we identified that Bach2 allows for improved persistence under *in vitro* exhaustion stress and *in vivo*. In addition, BACH2 OE allows for improved tumor control under a chronic exhaustion environment in human CART19-28ζ cells.

*SB* mutagenesis in DiSBey mice offers a powerful, complementary approach to CRISPR-Cas9 for identifying gene functions in engineered T cells. Unlike CRISPR, which typically induces homologous knockouts or OE of full-length genes, *SB* mutagenesis can introduce heterozygous GOF and LOF mutations, including truncated variants, within a single screen^16,43^. DiSBey also eliminates the need for transducing gRNA libraries *in vitro*, enabling spontaneous *in vivo* screenings. The Dox-inducible design allows mutagenesis in CD4⁺ and CD8⁺ T cells while minimizing leukemogenesis, and *in vivo* selection against deleterious mutations refines the mutant pool before screening. Furthermore, DiSBey’s design enables flexible screening across immune subsets by substituting alternative promoters to drive Cre recombinase expression.

Although Bach2 is known to prevent exhaustion under chronic virus infection^23–26^, its use in cancer-induced exhaustion for engineered T cells under immune-suppressive TME is unclear. We show that low ectopic Bach2 OE is responsible for improved persistence under chronic tumor antigenic stimulation while retaining the capacity to maintain effector phenotype. Supporting our CAR-T findings, a recent preprint reported that BACH2 OE in CD22-CD28ζ CAR-T cells induces transcriptional and functional features similar to 4-1BB-based CARs^44^. However, unlike our results, the therapeutic benefit in that model was limited to transient BACH2 expression during manufacturing^44^. These differences may stem from CAR design or chronic stimulation protocols. Together, both studies underscore the potential of modulating engineered T cell phenotypes by controlling ectopic BACH2 expression levels. Additionally, our identification of Bach2 under chronic TCR stimulation in CD8⁺ T cells suggests transgenic TCR therapies may similarly benefit from BACH2 OE.

In summary, our work demonstrates the novel application of controllable *SB* mutagenesis to identify enhanced T cell functionalities and highlights the OE of BACH2 as a potential candidate for improving CAR-T cell therapy. The DiSBey model and multiomic pipelines established here offer a foundation for broad functional interrogation of engineered T cells.

## Supplementary Figure Legends

**Supplementary Figure 1.**
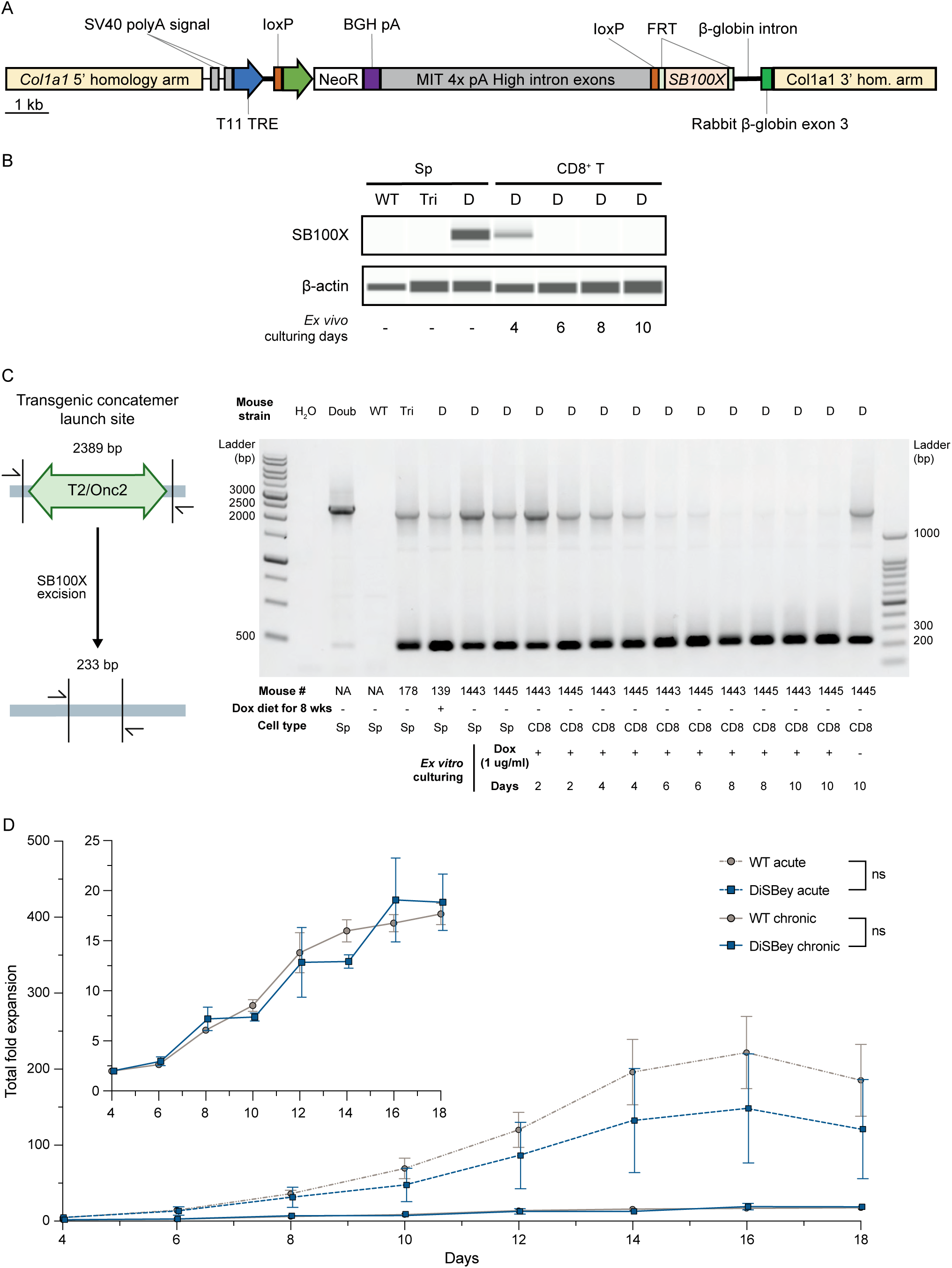
A) Schematic of the cassette sequences used to integrate T11-TRE-LSL-SB100X into mouse embryonic cells. B) Decay of SB100X expression in T cells of DiSBey mice in Dox-free culturing environment. CD8^+^ T cells from DiSBey mice (on Dox diet for at least 8 weeks) were cultured in culturing media without Dox for the indicated days. Splenocytes (“Sp”) from WT C57BL/6J, Triple transgenic (“Tri”), and DiSBey were controls for SB100X expression. C) Effective T2/Onc2 excision from the transgenic launching concatemers in Dox-treated CD8^+^ T cells from DiSBey mice *in vitro*. Splenocytes from double transgenic (“Doub”, no SB100X activity with T2/Onc2 and tetO-LSL-SB100X transgenes), Tri, and DiSBey mice were controls for T2/Onc2 excision. D) Fold expansion of WT and DiSBey mice under acute or chronic stimulation. n=4 from each group.

**Supplementary Figure 2.**
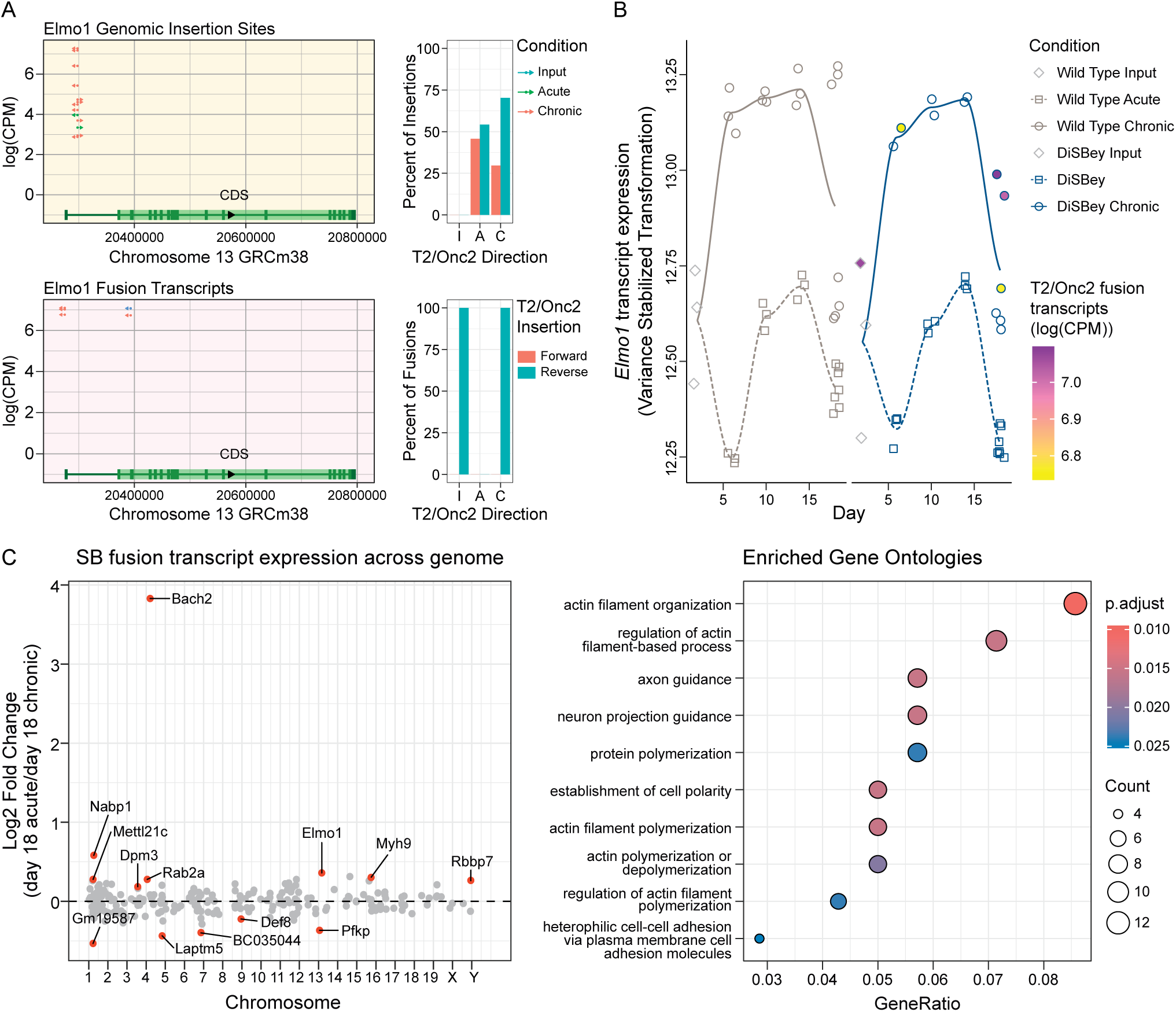
A) T2/Onc2 genomic and transcriptomic insertion diversity for *Elmo1*. B) Normalized expression of *Elmo1* from DESeq2 using the variance stabilizing transformation and compared against the counts per million T2/Onc2 fusion transcripts. C) Manhattan plot indicating fusion transcripts enrichment across the genome. D) GO pathway analysis on all differential transcripts or insertions of day 18 chronically stimulated DiSBey CD8^+^ T cells.

**Supplementary Figure 3.**
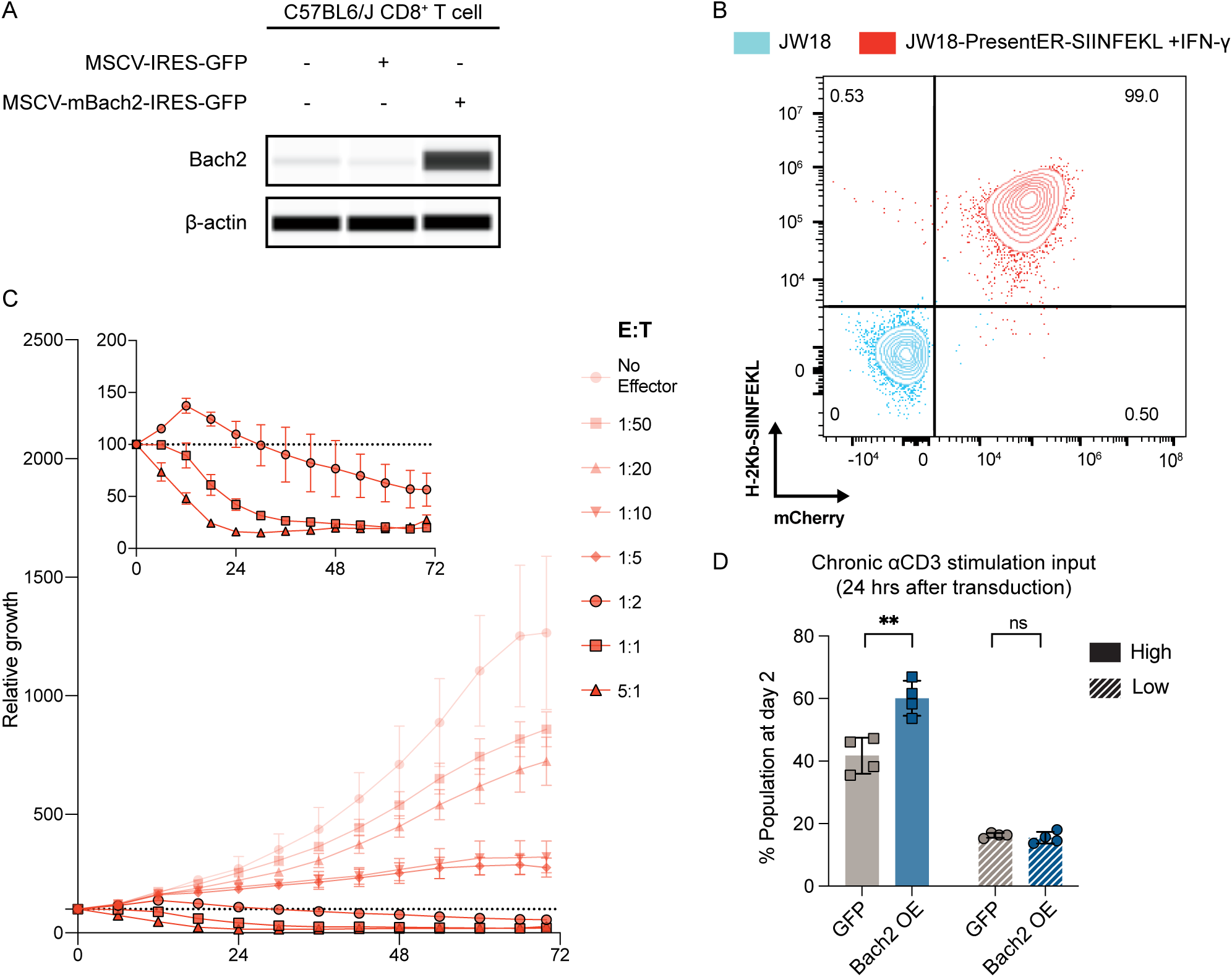
A) Bach2 expression of CD8^+^ T cells from C57BL/6J mice after Bach2 OE vector transduction. B) mCherry and H-2Kb-bound OT-I congenic peptide SIINFEKL presentation of JW18-SIINFEKL cells. C) Targetability of JW18-SIINFEKL cells against OT-I T cells at different E:T ratios. D) Population rate of Bach2 OE OT-I cells 24 hours after retroviral transduction. (Paired t-test, n=4 mice)

**Supplementary Figure 4.**
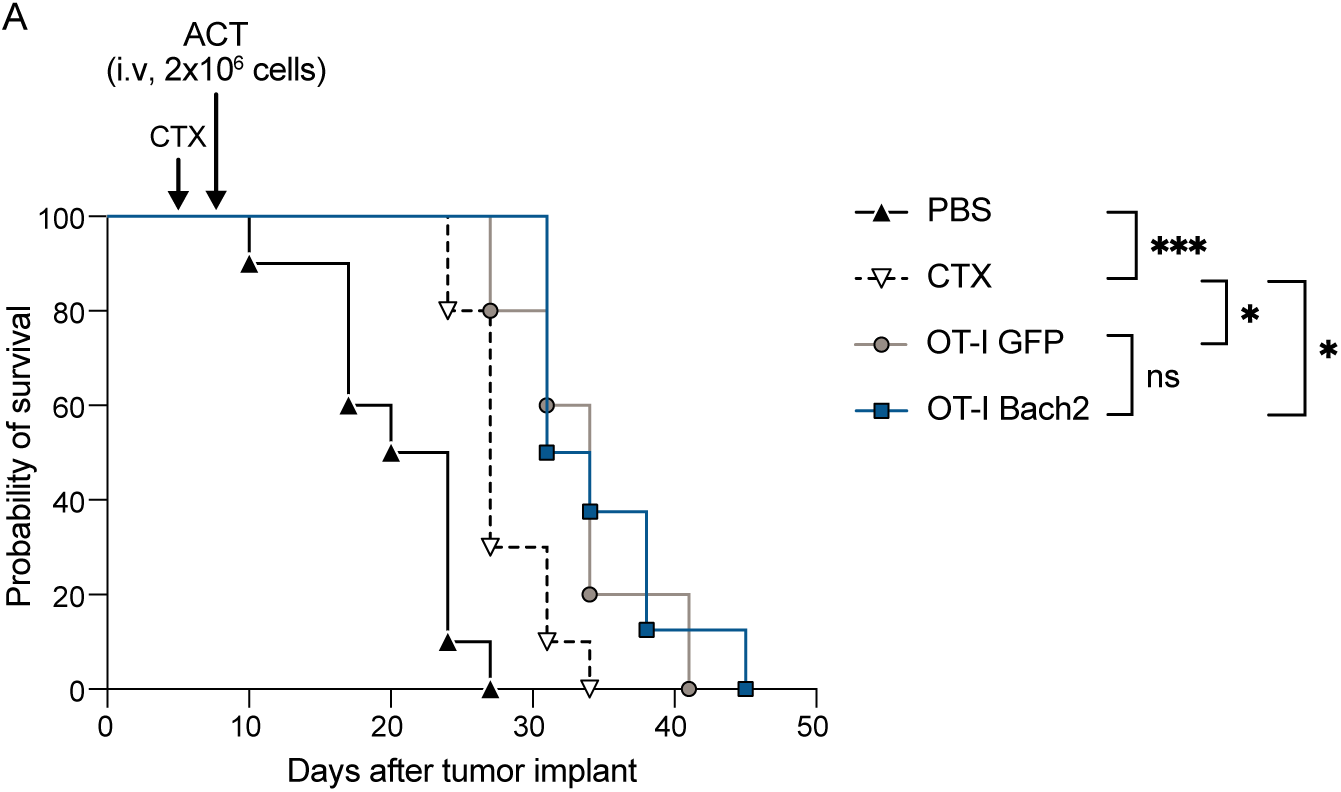
Survival of C57BL/6J mice transferred with OT-I GFP or Bach2 OE cells. (Log-rank (Mantel–Cox) test).

**Supplementary Figure 5.**
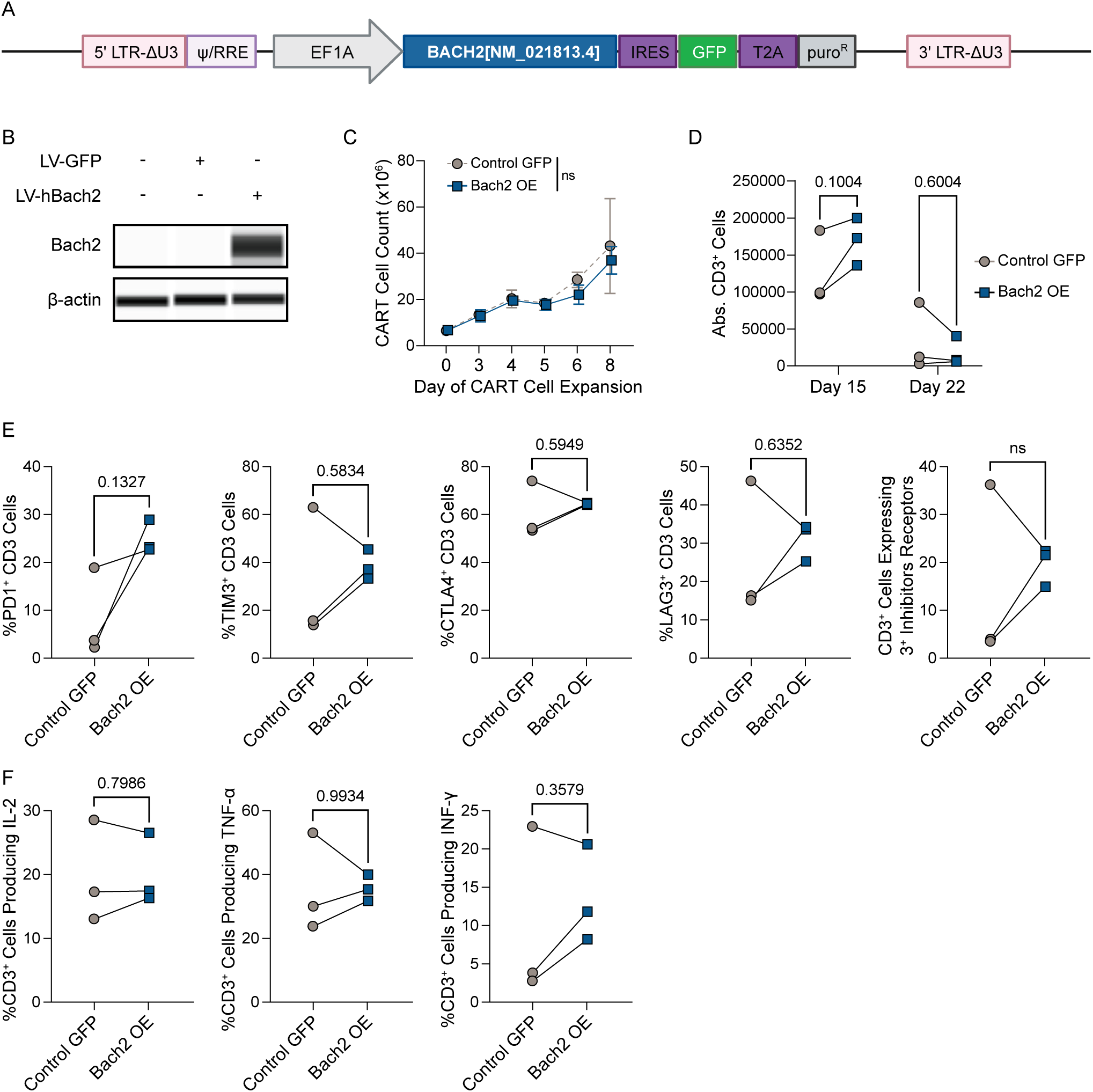
A) Vector schematic for lentiviral vector overexpressing human BACH2. B) Validation of BACH2 overexpression. C) Expansion of human CAR19-28ζ expressing with control or BACH2 OE vector during the first 8 days of CAR-T manufacturing. n=3 donors per group for panel 3-F. (Two-way ANOVA with Tukey‘s test) D) Proliferation of chronically exhausted CAR19-28ζ at indicated timepoints. (paired t-test) E) T cell inhibitory receptor expression of CAR19-28ζ cells at day 22 of the chronic exhaustion assay. Results were gated at CD3^+^ cells. (paired t-test) F) Pro-inflammatory cytokine expression of CAR19-28ζ cells at day 22 of the chronic exhaustion assay. Results were gated at CD3^+^ cells. (paired t-test)

**Supplementary Figure 6.**
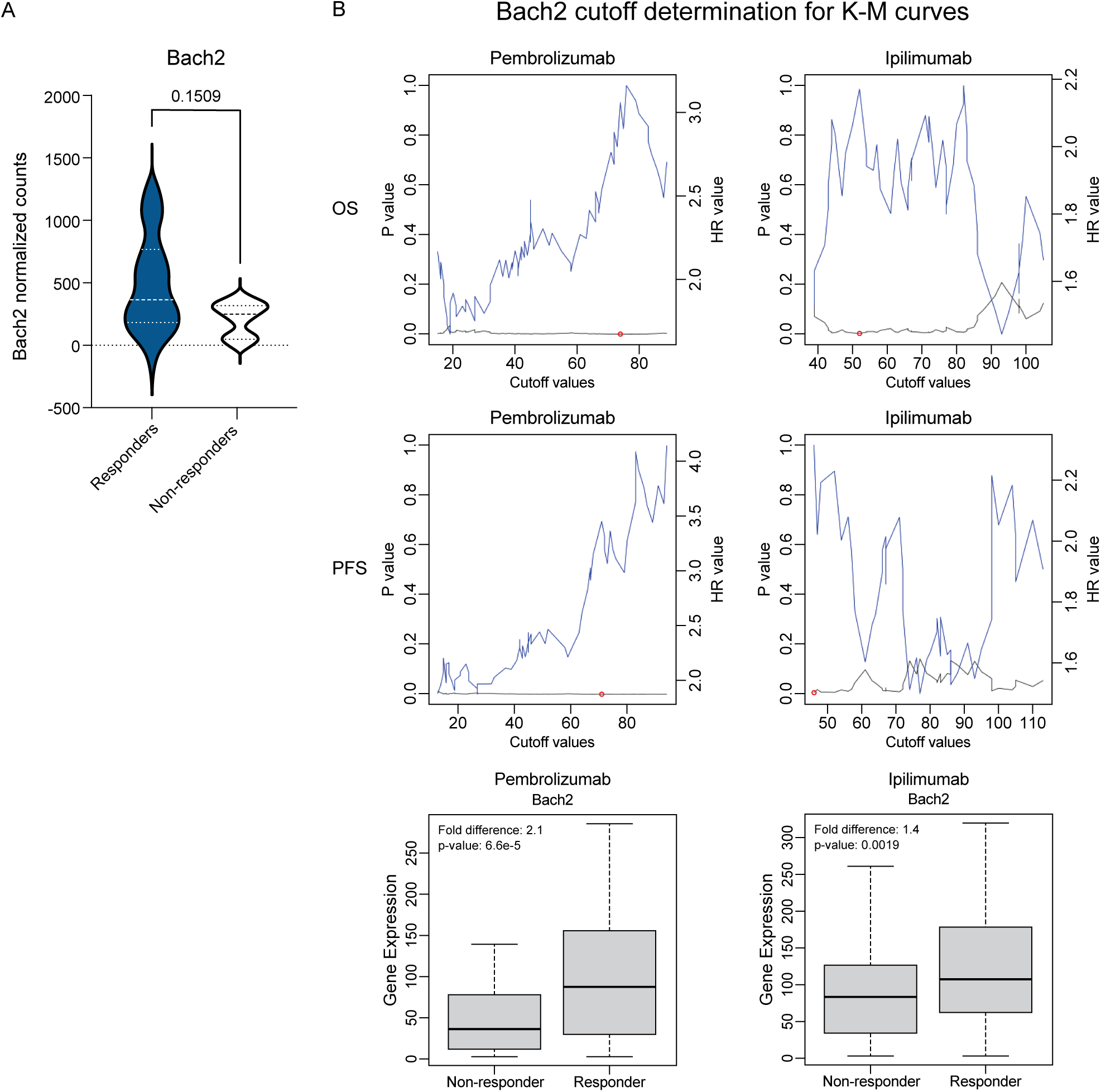
A) BACH2 expression comparing pre-infusion axi-cel products from responders and non-responders in the ZUMA-1 clinical trial (unpaired t-test, n=6 patients per group) Significance vs cutoff values between the lower and higher BACH2 expression for the survival analyses in a curated patient cohort treated with Pembrolizumab or Ipilimumab.

## Acknowledgements

We would like to thank members of the Genentech Genetically Engineered Mouse lab for mouse ES cell targeting and clone verification, the translational medicine team from Kite Pharma for sharing data related to the ZUMA-1 study, Dr. Scott W. Lowe for the Rosa-CAGS-LSL-rtTA3-IRES-mKate2 mouse, Dr. Matthew F. Mescher for OT-I/PL mouse, and Ms. Eva M. Baucom (Afferent Visual Science) for figure visualization. This work was funded by the American Cancer Society Research Professor Award #123939 to D.A.L., the Minnesota Partnership for Biotechnology and Medical Genomics to D.A.L., S.S.K., and I.M.S., Mayo Clinic Comprehensive Cancer Center to S.S.K., National Institutes of Health R37CA266344 to S.S.K., and an AIRP Grant from the University of Minnesota to I.M.S. and D.A.L.

Figure 1B, 2A, 4A, 4J, 5A, 6A, 7A, 7C, and Supplementary Figure 1C and 5A were created with BioRender.com.

