## Supplementary Methods for "*Sleeping Beauty* mutagenesis identifies *BACH2* and other regulators of CD8^+^ T cell exhaustion, persistence *in vivo*, and CAR-T function under tumor-associated chronic antigen stimulation"

*Cell line information*

The cancer cell lines are murine melanoma B16-F10 (ATCC CRL-6475), B16-Ova (Sigma-Aldrich SCC420), murine malignant peripheral nerve sheath tumor-like JW18[1], and human mantle cell lymphoma JeKo-1 (ATCC CRL-3006). All cell lines were routinely tested for the absence of mycoplasma contamination. To monitor OT-I cell killing, murine JW18 was transduced to express SIINFEKL peptide and mCherry (Addgene 102945), followed by puromycin selection (**Supplementary Figures 3B and 3C**). For evaluations of tumor killing by CAR-T cells, the JeKo-1 cells were transduced with luciferase-ZsGreen lentivirus (Addgene #39196) and sorted to 100% purity before experimentation.

*Mice information*

C57BL/6J, OT-I, and NOD-SCID-IL2rγ-/- (NSG) mice were purchased from Jackson Laboratories. OT-I/PL (Thy1.1$^+$ OT-I hereafter) was a gift from Dr. Matthew Mescher. Mice were housed at the University of Minnesota (IACUC protocols 1509-A33053, 2402-41779A, 2405-42060A, and 2405-42060A; IBC protocol 2205-40064H) or at the Mayo Clinic (IACUC protocol A00001767-16-R22; IBC protocol HIP00000252.43).

*Immunoblot analysis*

Cell pellets were placed on ice and lysed using RIPA cell lysis buffer (Sigma-Aldrich R0278) supplemented with protease- and phosphatase inhibitors (Sigma 11873580001 and P0044, respectively). Samples were vortexed at 2,000 RPM for 15 minutes at 4°C, then sonicated 20 times in one-second pulses. Lysates were then quantified using the Pierce BCA kit (ThermoScientific 23227). The 12-230 kDa size module (ProteinSimple SM-W004) of the Wes Simple Western (ProteinSimple) was used for immunoblotting analysis. Primary antibodies used at 1:50 for *SB* transposase (R&D Systems AF2798), 1:100 for BACH2 (Abcam ab243148), and

1:50 for beta-actin (Cell Signaling 8457). Secondary antibodies used at 1:50 for donkey anti-goat HRP (R&D Systems HAF109), 1:50 for goat anti-rat HRP (R&D Systems HAF005), and anti-rabbit HRP conjugate (ProteinSimple 042-206).

*Retrovirus production and tittering*

$7.5\times10^6$ low-passaged Plat-E (Cell Biolabs RV-101) cells were seeded in poly-L-lysine-coated 15 cm plates. The next day, plasmids containing MSCV control vectors (Addgene 20672 or 102796) or mBach2(NM_001109661.1)-IRES-GFP were transfected by TransIT®-LT1 Transfection Reagent (Mirus 2304) following manufacturer's protocol. One day later, replace media with media containing 1X ViralBoost Reagent (Alstem Cell VB100). MSCV-containing supernatant was collected 24 and 48 hours after ViralBoost Reagent addition, filtered with 0.45 μm filter, and concentrated with Retro-X™ Concentrator (TaKaRa 631456). Concentrated virus was flash frozen before storage in -80 °C until use.

*Murine T cell culturing*

Spleens were collected and passed through 70 μm strainers. Red blood cells were then lysed with 1X RBC Lysis Buffer (Invitrogen 00-4300-54) for 5 minutes before washing with FACS buffer. Naïve $CD8^+$ T cells were isolated  (Miltenyi 130-096-543) and resuspended at $1\times10^6$ cells per ml in mouse T cell media, which consisted with 10% FBS, 1% sodium pyruvate (MP Biomedicals 091682049), 1% Non-essential amino-acids (Gibco 11140050), 100U Pen/Strep (Corning 30002CI), and 50 nM of β-mercaptoethanol in RPMI and supplemented with 10 ng/mL of mouse IL-2 (R&D Systems 402-ML-020/CF). Naive $CD8^+$ T cells were then activated by CD3/CD28 Dynabeads (Gibco 11456D). 2 days later, magnetic Dynabeads were removed, and $CD8^+$ T cells were expanded in TCM for experiments. Fluorescent mKate2 positivity from DiSBey $CD8^+$ T cells was routinely checked with flow cytometry when applicable.

*Murine $CD8^+$ T cell transduction*

24 hours after CD3/CD28 Dynabeads activation, $CD8^+$ T cells were mixed with RV (MOI=5) before transferred into non-TC-treated plates coated with Retronectin (TaKaRa T100A). $CD8^+$ T cells were then spun at 2,000 g for 2 hours for spinoculation. Transduction efficiency was checked 24 hours after transduction with CytoFlex S for *in vitro* assays. For *in vivo* studies, transduced cells were sorted by BD FACSAria II before expansion.

*In vitro murine T cell assays*

To evaluate apoptosis, T cells were isolated at each indicated time point and stained with APC Annexin V (Tonbo Bio SKU 20-6409-T100) and 7-AAD (Tonbo Bio SKU 13-6993-T200) before being analyzed by flow cytometry. To compare proliferation activity of mouse $CD8^+$ T cells after chronic TCR stimulation, day 18 chronically stimulated T cells were stained with Violet Proliferation Dye 450 (BD 562158) following manufacturer's protocol and cultured for 96 hours before being analyzed by flow cytometry.

To assess OT-I cytotoxicity, $mCherry^+$ JW18-SIINFEKL cells were treated with mouse IFN-γ (100 ng/ml for 24 hours) before being co-cultured with chronically stimulated OT-I T cells at the indicated E:T ratio for 48 hours. Live cell imaging of the $mCherry^+$ area was recorded and analyzed by Incucyte SX5.

*In vitro competition assay with chronic tumor cell stimulation*

For target cells, irradiated (10 Gy) B16-F10 were treated with mouse IFN-γ (Cell Signaling 39127S, 100 ng/ml for 24 hours) to boost MHCI expression before pulsing with SIINFEKL peptide (100 nM for 2 hours). The unbound peptide was washed away with PBS before co-culturing with transduced OT-I. For effector cells, $CD8^+$ T cells from $Thy1.1^+$ OT-I mice were activated, transduced, and expanded as described previously. At day 4, OT-I expressing with Bach2-GFP and TagBFP from the same donor were mixed at 1:1 ratio. These OT-I mixture were then co-cultured with irr.B16-SIINFEKL (prepared as described above) at 1:1 E:T ratio. Co-culturing was repeated at day 8 and 12, and remaining OT-I cells were analyzed by flow cytometry at day 16.

*In vivo competition assay with B16-Ova model*

$5\times10^5$ B16-Ova were injected subcutaneously into the right flank of C57BL/6J mice (6-8 weeks old). 7 days later, mice were intravenously injected with $1\times10^6$ Thy1.1$^+$ OT-I T cells mixture, which contains Thy1.1$^+$ OT-I cells transduced with Bach2-GFP or TagBFP. At 7, 10, and 14 days after OT-I transfer, mice were euthanized for tissue collection, for which cardiac blood, right inguinal lymph nodes, spleen, and tumors were isolated for flow cytometry analysis.

*In vivo KPC-Ova model*

Kras$^{LSL\text{-}G12D/+}$;Trp53$^{LSL\text{-}R172H/+}$;p48-Cre (KPC) mice backcrossed to the C57BL6/J background[2] were bred to the ROSA26 LSL-EGFP-Ova strain generated by Strandt et al[3] to generate KPC-Ova mice. A cell line from primary KPC-Ova tumors was generated and used for the experiments. KPC-Ova cells were orthotopically implanted as previously described[4] to assess the relevance of Bach2 in vivo during antitumor immunity. Seven days after tumor implantation, mice received 120 mg/kg cyclophosphamide (Amneal Pharmaceuticals) i.p for lymphodepletion followed eight hours later by adoptive cell therapies (ACT) with $2\times10^6$ sorted OT-I Bach2 OE i.p. Mice then received 5 µg of IL-2 daily on days +1-3 after ACT. 10 days after ACT, pancreatic tumor-draining LNs and spleens were isolated and single cell suspensions were generated as previously described[4].

*Production of edited human CAR-T cells*

To generate CAR-T cells, human peripheral blood mononuclear cells (PBMCs) with SepMate PBMC isolation tubes (STEMCELL Technologies) from de-identified blood samples that were obtained under a Mayo Clinic IRB approved protocol. T cells from the PBMCs were isolated using

EasySep Human T Cell Isolation Kit with a Robosep machine (STEMCELL Technologies). The T cells were then activated with CD3/CD28 beads (Cell Therapy Systems Dynabeads™) at a 3:1 bead-to-cell ratio. 24 hours later, activated T cells were transduced at MOI=3 with lentiviral particles encoding a CD19-targeting CAR with a CD28ζ costimulatory domain as previously described[5]. The next day, CAR-T cells were transduced at MOI=3 with lentiviral particles generated with a BACH2 OE or control plasmid that contains a GFP and puromycin resistance sequence **(Supplementary Figure 5A)**. From day 3 to 8, CAR-T cells were maintained at $1x10^6$ cells/mL and supplemented the T-cell media with 1 µg/mL puromycin. On Day 6, CD3/CD28 beads were removed through magnetic separation and evaluated CAR expression and GFP positivity with flow cytometry. On day 8, the CAR-T cells were washed three times to remove puromycin before using them in experiments. During CAR-T cell expansion and *in vitro* studies, CAR-T cells were kept in T-cell media that is composed X-Vivo15 (Lonza) supplemented with 10% human serum albumin (Corning) and 1% penicillin-streptomycin-glutamine (Gibco).

*In vitro human CAR-T cell assays*

To evaluate changes in antigen-specific proliferation, T cells were co-cultured with JeKo-1 cells at a 1:1 ratio in a 96-well plate for 5 days. After 3 days, 100 µL of media was removed from all wells, and it was replaced with 100µL of fresh T cell media. On day 5, the absolute count of live $CD3^+$ cells was determined via a volumetric measurement with flow cytometry after staining samples with Zombie R718™ fixable viability kit and CD3 (clone OKT3) as described previously.

To evaluate changes in *in vitro* CAR-T cell cytotoxicity, we co-cultured CAR-T cells with luciferase+ JeKo-1 cells at the indicated E:T ratios. After 48 hours, killing was assessed with luminescence imaging on a GloMax Explorer (Promega) after adding 1µL D-luciferin (3mg/mL, Gold Biotechnology) per 100µL sample volume.

*Mantle cell lymphoma xenograft model*

NSG mice (6-8 weeks old females) were intravenously injected with $1x10^6$ luciferase$^+$ JeKo-1 tumor cells. 7 days later, mice were intraperitoneally injected with 3 mg luciferin, and luminescence imaging was performed with the IVIS to determine tumor burden for randomization. The next day, mice were randomized into groups based on tumor burden and intravenously injected with $1x10^6$ fresh day 22 CAR-T cells. Following CAR-T cell injection, the mice were regularly monitored for tumor burden with luminescence imaging and changes in weight.

*Nucleic acid isolation, esTag-seq, and RNA-seq*

Simultaneous genomic DNA and RNA of murine CD8$^+$ T cell samples were isolated using Quick-DNA/RNA MagBead (Zymo R2130). For esTAG-seq, genomic DNA was tagmented with UMI-loaded Tn5 transposase (seqWell 301230). For insertion site enrichment with PCR, primers targeting *SB* inverted regions (listed in **Supplementary Table 3**) were paired with the p5 forward primer in sequential nested PCR reactions. PCR products were purified with SPRI Beads and sequenced with Element Aviti 2x150 bp with a 20 nt i5 index read (to sequence the Tagify 10 nt i5 index and 10 nt UMI sequences). For bulk RNA-seq, mRNA was reverse-transcribed and sequenced at 2x150 bp.

*Sleeping Beauty sequencing data processing and analysis.*

Sequencing raw data was deposited at GSE302682 and GSE302683. T2/Onc2 fusions were quantified by trimming the RNA-seq data with trimmomatic then utilizing Arriba with a modified genome that included the T2/Onc2 construct as a chromosome. For RNA-seq expression quantification, reads included a T2/Onc2 construct were excluded, and the remaining reads were quantified with HISAT2 and Subread. T2/Onc2 fusions were quantified by removing the primer sequences and aligning the remaining reads to the genome using BWA MEM, deduplicated using UMI Tools, and quantified with a custom R script. Analysis pipeline manuscript is in preparation.

Initial fusion and insertion detection included only fusions or insertions with at least 3 deduplicated supporting reads. Additional filtering was applied downstream.

All downstream analyses were completed in R. The bottom 10% of expressed fusions were removed from each sample via quantile analysis. Differential analysis for both genomic insertions and fusion transcripts was performed at the gene level using DESeq2[34] with an offset of 1 count applied. Differential genomic insertions were identified as those with a p-value < 0.05 and with a log2 fold change > 0.5. Differential fusion transcripts were identified as those with a p-value < 0.05 and with a log2 fold change > 1. Overrepresentation analysis was conducted with clusterProfiler using Gene Ontologies (GO)[35] with a p-value of 0.01 and a q-value of 0.05.
